# Nutrient utilization and degradative enzyme activity of the dragon fruit canker pathogen, *Neoscytalidium dimidiatum*

**DOI:** 10.64898/2025.12.20.695650

**Authors:** Rachel E. Kalicharan, Romina Gazis, Jessie Fernandez

## Abstract

Dragon fruit canker (DFC), caused by the pathogenic fungus *Neoscytalidium dimidiatum*, is a severe disease that threatens dragon fruit production worldwide. Current management efforts largely rely on fungicide applications and sanitation measures; however, the pathogen’s molecular genetic characterization, nutrient utilization preferences, and extracellular enzyme activities remain poorly understood. Using calcofluor white-based fluorescence microscopy, we demonstrate that nutrient limitation significantly restricts the growth and maturation of *N. dimidiatum*, impairing melanization and sporulation. Carbon utilization assays revealed a preference for maltose, suggesting reliance on starch-derived sugars. In addition, nitrogen utilization assays indicated efficient assimilation of complex organic nitrogen sources rich in peptides and amino acids. Finally, we experimentally validated extracellular enzymatic activities involved in host macromolecule degradation, including cellulase, amylase, pectinase, and protease activities. Collectively, these findings provide new insights into the physiology and pathogenic potential of *N. dimidiatum* and establish a foundation for developing improved strategies to mitigate DFC.

**Impact Statement:** Neoscytalidium *dimidiatum* is the causal agent of dragon fruit canker (DFC), one of the most devastating diseases affecting dragon fruit. Here, we investigated key features of N. dimidiatum biology relevant to disease development, including nutrient preferences and degradative enzyme activity. Our findings indicate that *N. dimidiatum* preferentially utilizes maltose as a carbon source, consistent with the use of starch-derived sugars, and grows efficiently on rich, complex organic nitrogen sources. Plate-based enzymatic assays aligned with these nutrient utilization patterns, revealing cellulase, amylase, pectinase, and protease activities. Collectively, this study provides new insights into *N. dimidiatum* physiology and establishes a foundation for future virulence studies that can ultimately support the development of improved DFC mitigation strategies.

## Introduction

Dragon fruit (*Selenicereus* spp., syn. *Hylocereus* spp.), commonly known as pitaya or pithaya, is an important perennial tropical crop cultivated worldwide, including Mexico, Central and South America, the Caribbean, South Africa, Asia, and Australia (1). With rising global demand, the dragon fruit industry has grown into a billion-dollar market, producing millions of metric tons annually (2). Despite this rapid expansion, production is increasingly jeopardized by several factors, most notably dragon fruit canker (DFC) caused by the fungal pathogen *Neoscytalidium dimidiatum*. This pathogen is notorious for its broad host range, causing dermatological infections in humans and diseases in woody and herbaceous plants such as almond, citrus, grapevine, and dragon fruit (3–8). DFC is considered the most destructive disease of dragon fruit globally, leading to severe economic losses reaching 80% in some cases (5). Infection occurs in both the vegetative tissues and fruit, primarily through natural openings such as stomata or mechanical wounds (9), although *N. dimidiatum* can also form appressoria to mechanically penetrate into the host epidermis (10). Symptoms begin as sunken, orange chlorotic spots on cladodes, progressing to mesophyll collapse and necrosis, which produce reddish-brown lesions that serve as sites for pycnidia development (10, 11). These pycnidia are where conidia (asexual spores) are formed, and as the lesions dry, characteristic “shot-holes” form on the cladodes (10, 11).

Given its aggressive nature, most research has focused on practical management strategies, including early detection, fungicide application, and sanitation practices (5, 11, 12). Recently, molecular studies have begun to elucidate the basis of *N. dimidiatum* pathogenicity, including whole-genome sequencing, effector identification, host defense gene expression, metabolite profiling, and biocontrol exploration (9, 13, 14). However, critical aspects, such as nutrient preference/utilization and validation of host-degradative enzymes, remain poorly understood. Here, we provide experimental evidence of *N. dimidiatum* carbon and nitrogen utilization preferences and demonstrate the activity of several fundamental degradative enzymes. We show that nutrient limitation significantly affects the pathogen’s growth, melanization, and sporulation. Carbon utilization assays reveal a preference for disaccharides, such as maltose, while nitrogen assays indicate efficient assimilation of complex, peptide-rich sources like yeast extract. Furthermore, *in vitro* plate-based enzymatic assays confirm the presence of amylase, cellulase, pectinase, and protease. Taken together, these findings offer key insights into the metabolic profile and virulence mechanisms employed by *N. dimidiatum* to exploit host resources, compromise host physiology, and drive the development of DFC symptoms.

## Methods

### Fungal Strains

The fungal strain *N. dimidiatum* was provided by Dr. Romina Gazis (University of Florida). Cultures were maintained on potato dextrose agar (PDA; 39 g/L; IBI Scientific) and incubated at 28 °C in the dark.

### Carbon and Nitrogen Source Testing

To assess carbon and nitrogen preferences, 4 mm agar plugs of 7-day-old PDA cultures were inoculated onto modified Czapek-Dox medium (1 g K_2_HPO_4_, 0.5 g MgSO_4_ • 7H_2_O, 0.5 g KCl, 0.01 g FeSO_4_ • 7H_2_O, distilled water to 1 L, pH adjusted 6.5 with HCl and solidified with 15 g agar before autoclaving 121 °C for 20 min). Media were supplemented with either 1% (w/v) carbon or 0.1% (w/v) nitrogen. Plates were incubated at 28 °C in the dark. Colony morphology and pigmentation were documented at 3, 5, and 7 days post-inoculation (dpi), and conidial production was quantified. Plates were imaged with an Epson Workforce Scanner.

### Sporulation assay

Spores were harvested from PDA cultures at 3, 5, or 7 dpi, suspended in water, and filtered through 3 layers of Miracloth. Suspensions were centrifuged at 3900 rpm for 5 min, supernatant discarded, and pellets resuspended in 1 mL of water. Spore counts were performed using a hemocytometer in triplicate (10 µL per count).

### Heat map generation and scoring criteria for melanization and sporulation

Melanization and sporulation were scored on a four-point scale (0-3) and visualized as heatmaps. Melanization scores were defined using reference phenotypes observed in Figure 1: 0, minimal colony growth with no/trace melanization; 1, full plate coverage with little melanization; 2, full coverage with moderate melanization; and 3, full coverage with strong melanization. Sporulation scores were assigned based on conidial concentration ranges: 0, ≤5 × 10^4^ spores/mL; 1, 5.01 × 10^4^–5 × 10^5^ spores/mL; 2, 5.01 × 10^5^–5 × 10^6^ spores/mL; and 3, ≥5.01 × 10^6^ spores/mL. Scores were mapped to the corresponding color for visualization.

**Figure 1.**
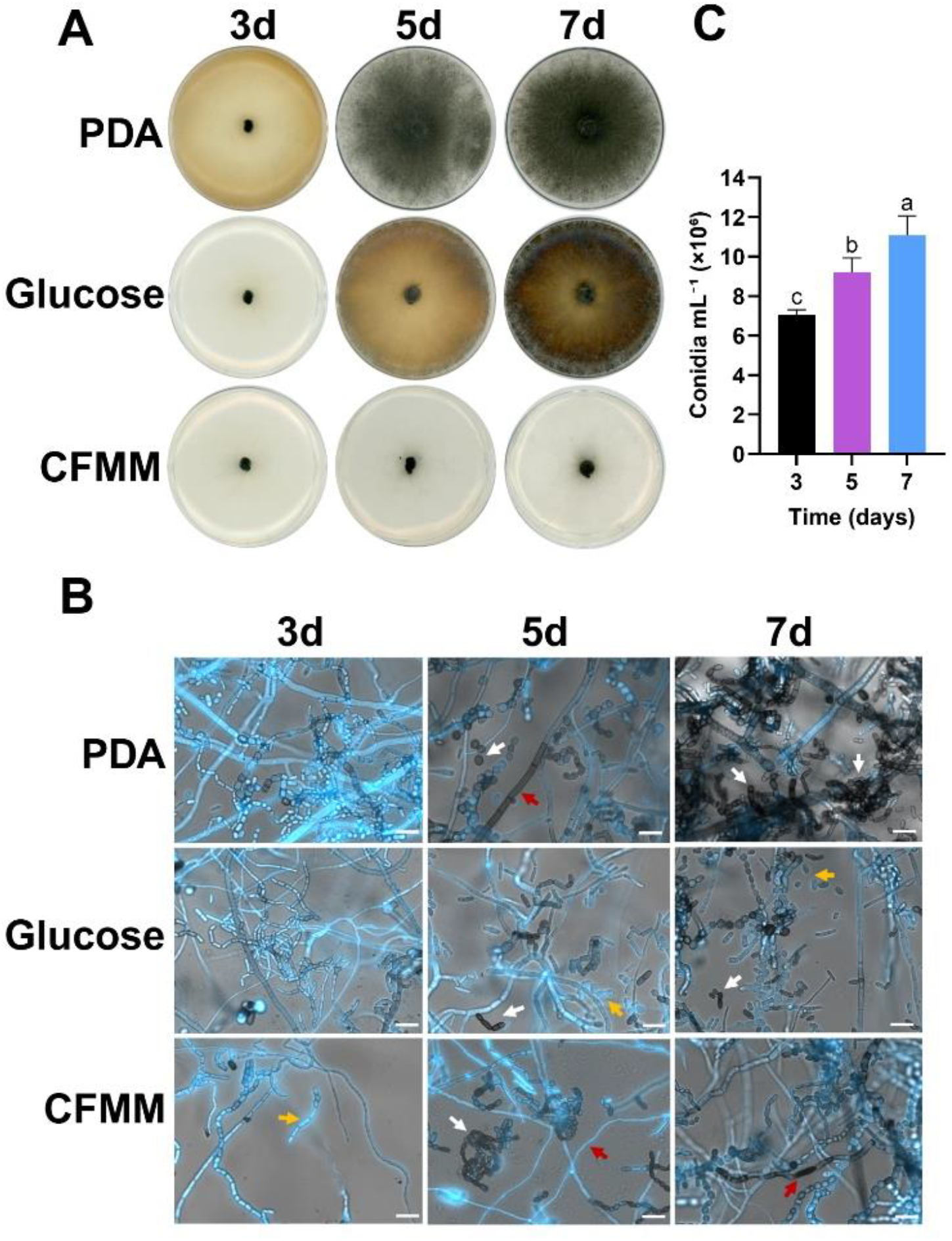
Development and melanization of *Neoscytalidium dimidiatum* under various nutrient conditions. *N. dimidiatum* was grown on microscope slides containing Potato Dextrose Agar (PDA), modified Czapek-Dox minimal media (Carbon-Free minimal media, CFMM), or CFMM supplemented with 1% glucose. Slides were examined and stained with calcofluor white at 3 days post-inoculation (dpi), 5 dpi, and 7 dpi. **A**. Colony morphology of *N. dimidiatum* grown on PDA, CFMM, or CFMM supplemented with 1% glucose at 3, 5, and 7 days dpi. **B**. Calcofluor white fluorescence microscopy of *N. dimidiatum* grown on PDA, CFMM, and CFMM + 1% w/v glucose. At 3 dpi, samples showed little to no melanization across conditions, as indicated by strong calcofluor white fluorescence; sporulation was most evident on PDA. At 5 dpi, melanization became apparent, particularly in PDA (darkened hyphae and spores) and in CFMM (darkened spores). Reduced calcofluor white signal in darkened structures is consistent with decreased dye penetration in melanized cell walls. At 7 dpi, PDA-grown cultures showed extensive melanization and abundant sporulation, whereas cultures grown on CFMM and CFMM + glucose exhibited delayed melanization, reflected by higher calcofluor white uptake and fluorescence. White arrows indicate melanized spores, orange arrows indicate unmelanized spores, and red arrows indicate hyphae. Scale bar = 20 µm. **C**. Conidial concentration of *N. dimidiatum* grown on PDA at 3, 5, and 7 dpi. One-way ANOVA with Tukey’s multiple-comparisons test; *p* < 0.05; *n* = 3. Created in https://BioRender.com

### Enzymatic Activity Assays

4 mm agar plugs of 7-day-old *N. dimidiatum* were cultured on modified Czapek-Dox medium supplemented with 1% w/v carboxymethylcellulose sodium salt (cellulase; TCI), 1% w/v starch (amylase), 1% w/v citrus pectin (pectinase; Sigma), or 10% w/v skim milk (protease). Plates were incubated at 28 °C for 3 days before evaluating activity. For cellulase activity, plates were stained with 0.1% Congo Red solution for 15 minutes, then cleared with three 5-minute washes of 1 M NaCl, and zones of clearance were assessed. For amylase and pectinase activity, plates were stained with Gram’s Iodine for 5 minutes (amylase) or 10 minutes (pectinase), excess iodine was poured off, and starch/pectin degradation was assessed by the intensity of staining and the presence of zones of clearance. For protease activity, a zone of clearance from the inoculation site was assessed.

### Fungal Cell Microscopy

Microscopic analysis of fungal development was performed on PDA, carbon-free minimal medium (CFMM), and CFMM supplemented with 1% w/v glucose. Media were applied to autoclaved microscope slides and dried prior to inoculation with 10 µL of a spore suspension (1 × 10^4^ spores/mL). Slides were incubated at 28 °C in humid chambers and imaged at 3, 5, and 7 dpi using a Zeiss Axio Observer 7 microscope. For calcofluor white staining, samples were treated with one drop of calcofluor white (Sigma) and one drop of 10% KOH per 3 cm of slide, incubated for 1 min, and imaged. Images were processed using Zen Microscopy Software v3.13 (Zeiss).

### Statistical Analysis

Statistical analyses were performed using a one-way ANOVA and a two-way ANOVA, with Dunnett post hoc and Tukey post hoc tests, comparing each treatment to the control. All experiments included at least three biological replicates, each with three replicates. Differences were considered statistically significant at *p* < 0.05.

## Results

### Nutrient deficiency impacts melanization and sporulation of *N. dimidiatum*

To assess how nutrient availability influences the growth, development, and maturation (conidiation and melanization) of *N. dimidiatum*, cultures were established on potato dextrose agar (PDA), carbon-free minimal medium (CFMM), and CFMM supplemented with 1% glucose. PDA served as a nutrient-rich control, while CFMM and glucose-supplemented CFMM represented nutrient-limited conditions. Fungal development was monitored microscopically at 3, 5, and 7 days post-inoculation (dpi), corresponding to key stages of pigmentation (melanization). At 3 dpi, colonies on PDA exhibited extensive mycelial expansion but remained white, indicating early developmental stages. Similar vegetative growth was observed on CFMM and glucose-supplemented CFMM, with all cultures appearing predominantly white (Figure 1A). Samples were stained with calcofluor white (CFW), which binds to chitin and β-glucans in the fungal cell wall and fluoresces upon binding (Figure 1B) (15). Fluorescence intensity was comparable across all conditions, consistent with the presence of non-melanized hyphae and spores.

By 5 dpi, clear differences in maturation emerged, and were largely complete by 7 dpi under nutrient-rich conditions (Figure 1A). PDA-grown samples displayed pronounced melanization in both hyphae and spores, whereas CFMM-grown cultures exhibited melanization primarily in spores, with hyphae remaining largely non-melanized. Glucose-supplemented CFMM supported growth but showed minimal melanization in either hyphae or spores (Figure 1B). Correspondingly, PDA-grown samples and melanized spores in CFMM exhibited reduced CFW fluorescence compared to glucose-grown samples, reflecting decreased dye accessibility in melanized cell walls (15). Sporulation was also markedly reduced under CFMM and glucose conditions compared to PDA (Figure 1B).

At 7 dpi, these developmental differences became more pronounced. PDA-grown cultures were fully melanized, forming dense hyphal networks and abundant spores. In contrast, CFMM and glucose-grown cultures remained similar to their 5 dpi state, with melanization restricted to spores but not hyphae, stronger overall CFW fluorescence, and reduced sporulation compared to PDA (Figure 1B). Taken together, these observations indicate that nutrient limitation significantly impairs *N. dimidiatum* maturation, particularly affecting melanization and sporulation.

### *N. dimidiatum* exhibits carbon-source preferences for maltose and glucose

To establish a baseline for normal development under nutrient-rich conditions, *N. dimidiatum* was first cultured on PDA, where it exhibited robust growth, progressive melanization, and abundant sporulation over time (Figure 1C). This phenotype served as a reference for evaluating how nutrient limitation and defined carbon sources influence colony expansion and key maturation markers, including pigmentation (melanization) and conidiation.

Following our initial assessment, we evaluated the effect of individual carbon sources on fungal development. *N. dimidiatum* was grown on CFMM alone and CFMM supplemented with various carbon sources, including monosaccharides (glucose, galactose), polyols (glycerol, mannitol, sorbitol), disaccharides (sucrose, maltose, lactose), and the polysaccharide starch. Cultures were incubated at 28 °C in the dark and evaluated at 3, 5, and 7 dpi for colony morphology, melanization, and conidial production.

At 3 dpi, all treatments exhibited similar early-stage growth, covering only a portion of the plate with no visible melanization (Figure 2A). By 5 dpi, clear differences arose: glucose, sucrose, and maltose supported faster colony expansion and earlier melanization, indicating accelerated maturation compared to the other carbon sources. At 7 dpi, glucose- and maltose-supplemented cultures displayed the most advanced development, with extensive melanization and dense colonies.

**Figure 2.**
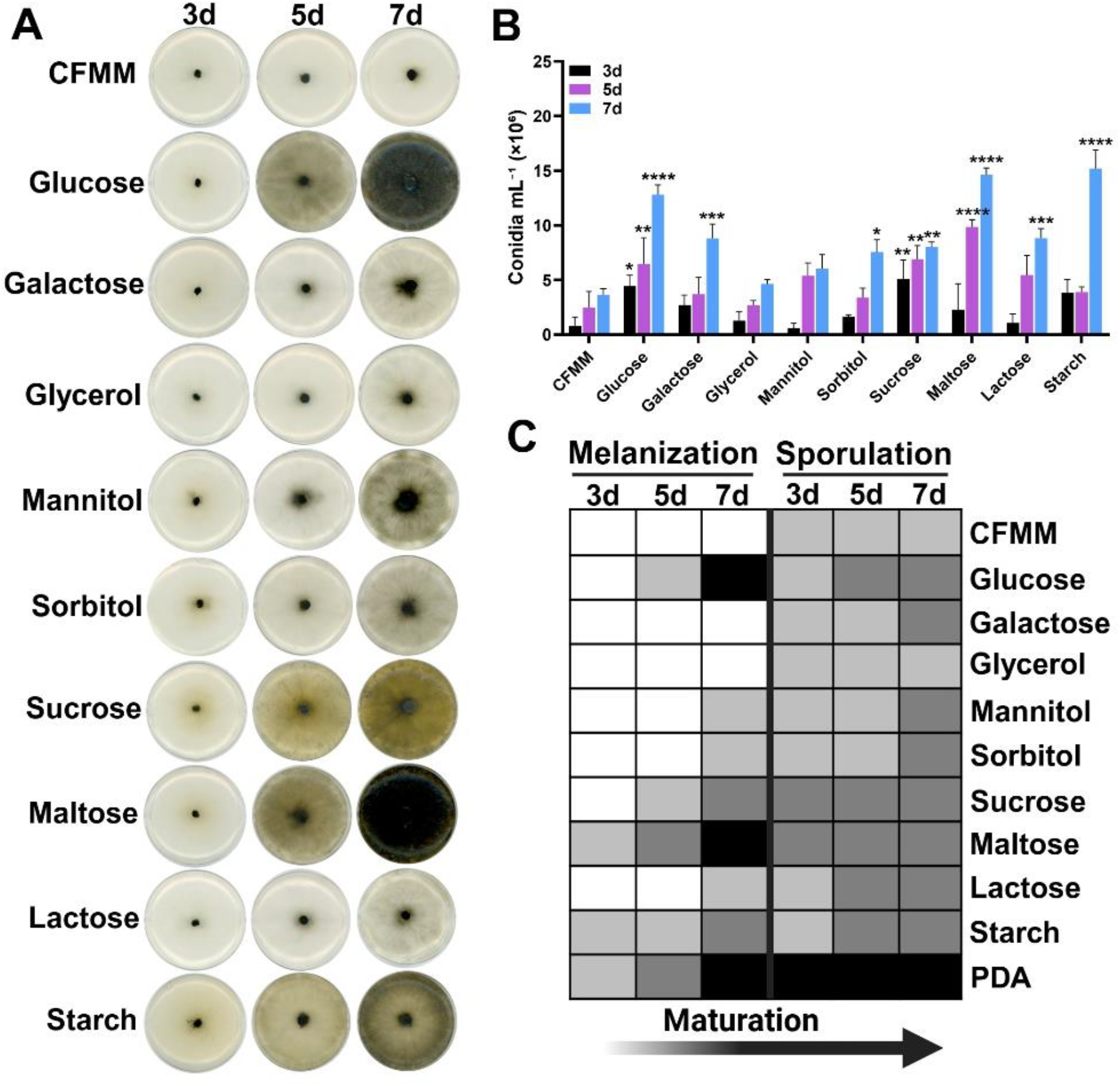
*N. dimidiatum* Carbon Source Utilization. **A**. Representative colony morphology of *N. dimidiatum* grown on modified Czapek-Dox minimal medium supplemented with 1% w/v carbon source at 3, 5, and 7 days post-inoculation (dpi). **B**. Quantification of conidial production under each condition at 3, 5, and 7 dpi. Statistical significance was determined by two-way ANOVA with Dunnett’s multiple-comparisons test. * indicates *p* < 0.05, ** indicates *p* < 0.01, *** indicates *p* < 0.001, **** indicates *p* < 0.0001. **C**. Composite heat map summarizing melanization and sporulation scores across carbon sources. Darker shading indicates greater melanization and/or higher sporulation reflecting advanced developmental progression. Created in https://BioRender.com

Conidial quantification revealed that all carbon sources supported growth, but sporulation varied significantly among treatments. Most supplemented media produced denser colonies and higher conidia counts than CFMM alone, with several carbon sources showing significantly increased sporulation (*p* < 0.05; Figure 2B). A composite heat map summarizing growth, melanization, and sporulation (Figure 2C) highlighted maltose and glucose as the most favorable carbon sources, promoting robust colony development, strong pigmentation, and high conidial output. Other carbon sources, such as sucrose and starch, supported either melanization or sporulation, but not both consistently. Notably, glycerol enhanced sporulation relative to CFMM but yielded weak melanization and poor overall growth. Overall, these findings indicate that maltose and, to a lesser extent, glucose provide optimal carbon conditions for *N. dimidiatum*, jointly promoting vigorous growth, melanization, and sporulation.

### *N. dimidiatum* demonstrates high affinity for complex, peptide-rich nitrogen sources

In addition to carbon preference, we assessed nitrogen utilization, as nitrogen availability is a key determinant of fungal fitness and pathogenicity (16). *N. dimidiatum* was cultured on nitrogen-free minimal medium (NFMM) or NFMM supplemented with individual nitrogen sources, including inorganic forms (ammonium chloride [NH_4_Cl], ammonium sulfate [(NH_4_)_2_SO_4_], sodium nitrate [NaNO_3_], potassium nitrate [KNO_3_]), urea, small amino acids (L-Asparagine), polypeptides (peptone, casein hydrolysate), and the complex digest yeast extract.

At 3 dpi, colonies grown on peptone, casein hydrolysate, and yeast extract exhibited extensive growth, covering most of the plate surface (Figure 3A). By 5 dpi, robust growth persisted in the casein hydrolysate and yeast extract treatments, accompanied by increased melanization in several other nitrogen sources, particularly ammonium salts, nitrates, and urea. At 7 dpi, cultures grown on yeast extract and casein hydrolysate remained the most visually similar, displaying dense mycelial growth and pronounced melanization. Although maturation was evident across multiple nitrogen sources, colonies grown on NH_4_Cl, (NH_4_)_2_SO_4_, L-Asparagine, urea, and peptone exhibited thin peripheral mycelial regions and visible underlying media, suggesting suboptimal growth (Figure 3A). These conditions also produced localized white patches, likely indicating physiological stress associated with inefficient nitrogen assimilation (17). Sporulation analysis revealed that yeast extract most strongly supported both growth and conidiation. At 3 dpi, all other nitrogen sources resulted in significantly reduced sporulation compared to yeast extract, with differences becoming more pronounced at 5 and 7 dpi (Figure 3B). These findings demonstrate that yeast extract is the most favorable nitrogen source, promoting coordinated fungal development and maturation (Figure 3C).

**Figure 3.**
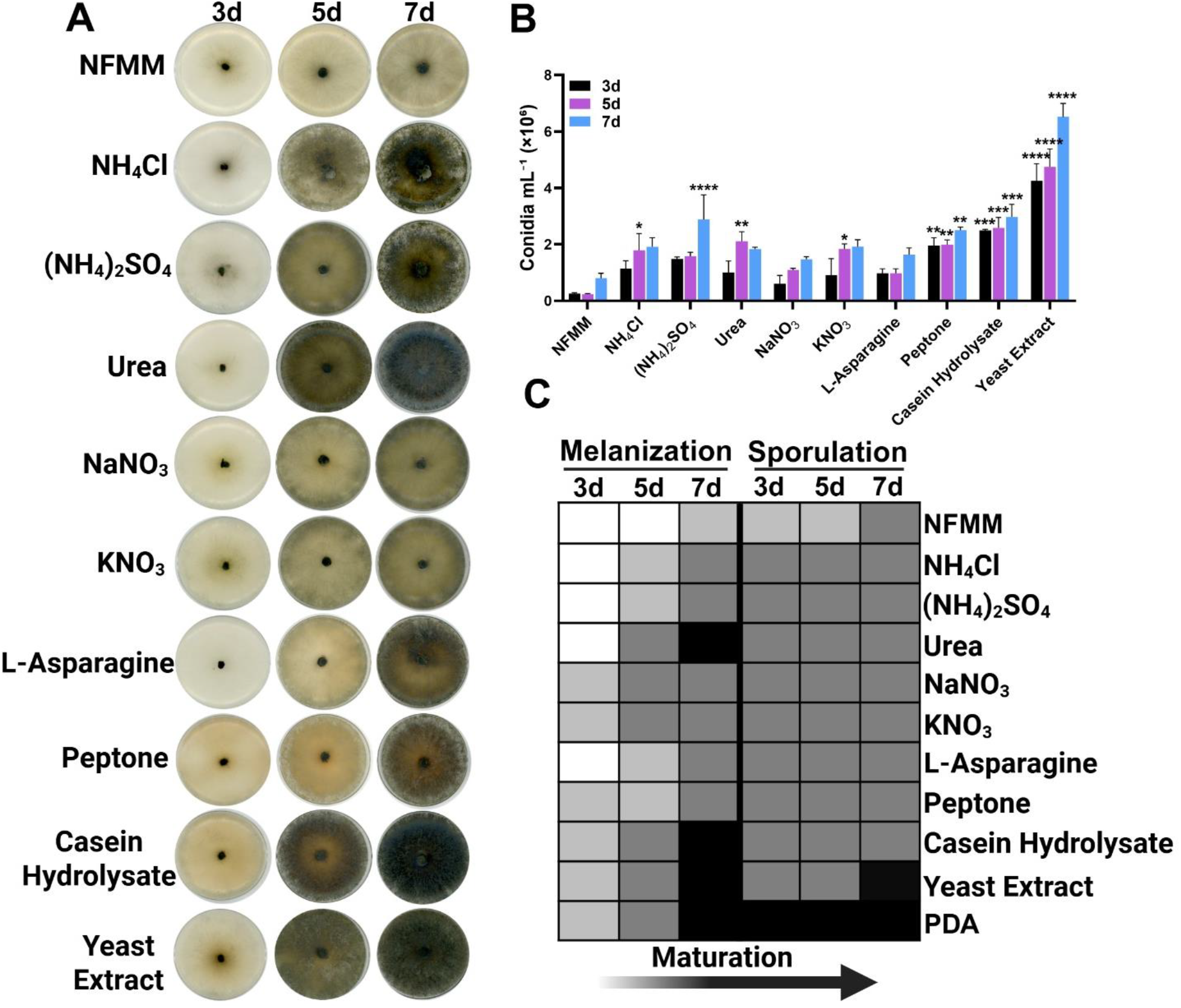
*N. dimidiatum* Nitrogen Source Utilization. **A**. Representative colony morphology of *N. dimidiatum* grown on modified Czapek-Dox minimal media, supplemented with 0.1% w/v nitrogen at 3, 5, and 7 dpi. **B**. Quantification of conidial production under each condition at 3, 5, and 7 dpi. Statistical significance was determined using a two-way ANOVA with Dunnett post-hoc, with one asterisk (*) indicating *p* < 0.05, two asterisks (**) indicating *p* < 0.01, three asterisks (***) indicating *p* < 0.001, and four asterisks (****) indicating *p* < 0.0001. **C**. Composite heat map summarizing melanization and sporulation scores across nitrogen sources. Darker colors indicate greater melanization and/or higher sporulation, reflecting advanced developmental progression/maturation. Created in https://BioRender.com

### *N. dimidatum* possesses plant cell wall-degrading enzymatic activity

To assess the secretion of plant cell wall-degrading enzymes (PCWDEs) and proteases by *N. dimidiatum*, we performed *in vitro* plate-based substrate degradation assays. Genomic analyses predict that *N. dimidiatum* encodes a diverse repertoire of carbohydrate-active enzymes (CAZymes) and PCWDEs (9). Combined with our observations of fungal growth on complex carbohydrates and peptide-rich nitrogen sources, we tested extracellular activities of cellulase, amylase, pectinase, and protease at 3 dpi.

Minimal medium was supplemented with 1% carboxymethylcellulose (CMC; cellulase), starch (amylase), pectin (pectinase), or 10% skim milk (protease). Plates were inoculated with fungal plugs, while uninoculated substrate plates and inoculated minimal medium plates processed with identical stains were included as negative controls. On CMC-amended plates, Congo Red staining followed by NaCl washing revealed a distinct clearance zone surrounding the inoculum (Figure 4A), consistent with cellulase-mediated CMC degradation (18). No clearing was observed in control plates, confirming substrate-dependent activity.

**Figure 4.**
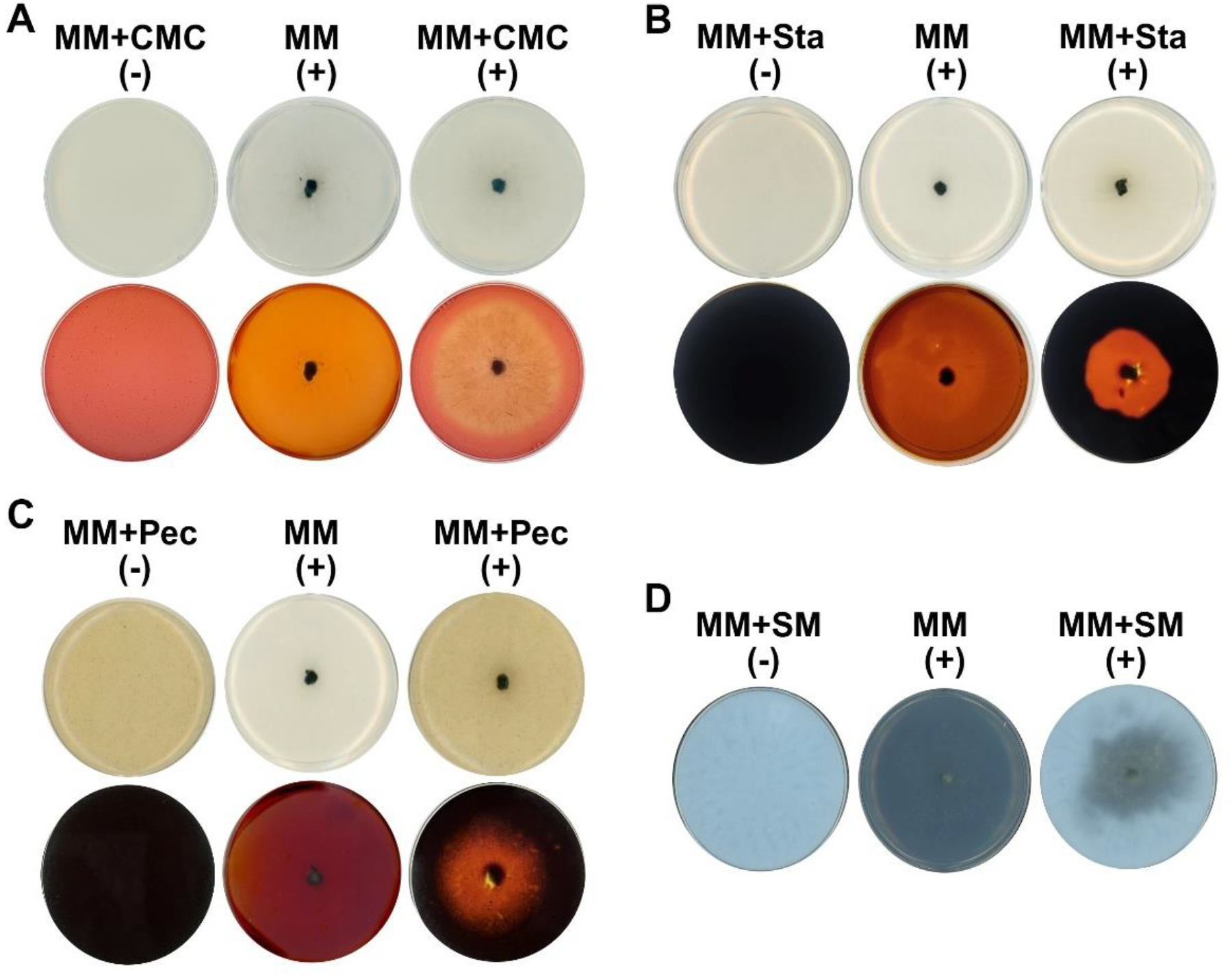
Extracellular Enzymatic Activity of *N. dimidiatum*. *N. dimidiatum* was cultured on modified Czapek-Dox minimal medium (MM) containing either 1% (w/v) carboxymethyl cellulose (CMC), starch, or pectin, or 10% (w/v) skim milk (SM), and imaged at 3 days post-inoculation (dpi). For each substrate, plates included MM + substrate (−) (uninoculated control), MM (+) (inoculated MM control), and MM + substrate (+) (inoculated test). **A**. Cellulase activity on CMC-amended plates. A zone of clearance on MM + CMC (+) relative to MM + CMC (−) indicates cellulase activity. **B**. Amylase activity on starch-amended plates. Reduced staining on MM + Sta (+) relative to MM + Sta (−) indicates starch hydrolysis. **C**. Pectinase activity on pectin-amended plates. Reduced staining on MM + Pec (+) relative to MM + Pec (−) indicates pectin degradation. D. Protease activity on skim milk agar. A clear halo on MM + SM (+) relative to MM + SM (−) indicates protease activity. Created in https://BioRender.com

Amylase activity was assessed on starch-supplemented plates stained with Gram’s iodine, which binds intact starch polymers (19). A strong clearance halo was observed around the fungal plug (Figure 4B), indicating starch degradation, in accordance with moderate growth and sporulation on starch-supplemented media in carbon utilization assays (Figure 2A-C). Similarly, pectin-supplemented plates stained with iodine exhibited localized clearing surrounding the inoculation site (Figure 4C), supporting pectinase activity.

Protease activity was confirmed on skim milk agar, where fungal growth produced a clear zone surrounding the inoculum (Figure 4D), indicating casein degradation. This aligns with the nitrogen utilization assays, where *N. dimidiatum* grew robustly on casein hydrolysate and yeast extract (Figure 3A-C). Collectively, these results demonstrate that *N. dimidiatum* secretes extracellular enzymes capable of degrading cellulose, starch, pectin, and protein substrates under *in vitro* conditions (Figure 4A-D), consistent with its functional capacity for exploiting plant-associated polysaccharides and protein-derived nitrogen sources.

## Discussion

Dragon fruit canker (DFC), caused by *Neoscytalidium dimidiatum*, is a highly destructive disease that threatens global dragon fruit production. Effective management of DFC requires a deeper understanding of pathogen biology, particularly how nutrient acquisition and metabolic flexibility influence fungal fitness and pathogenicity. This study provides the first experimental characterization of carbon and nitrogen utilization preferences in *N. dimidiatum* and validates the secretion of extracellular enzymes associated with plant tissue degradation, establishing a physiological framework for understanding its pathogenic lifestyle.

Under nutrient-limited conditions, microscopic examination revealed that restricted nutrient availability considerably delays key developmental processes, most notably melanization and sporulation. Both traits are widely recognized as indicators of fungal fitness and virulence in many plant-pathogenic fungi, including *N. dimidiatum* (15, 17, 20). These effects were most pronounced at 5 and 7 dpi, when melanization and sporulation were markedly reduced compared to PDA-grown cultures. Impaired melanization has critical consequences for fungal fitness, as melanin confers protection against oxidative stress, antimicrobial compounds, and metal toxicity, and is essential for infection-related structures in diverse fungal pathogens (20, 21). Similarly, reduced sporulation under nutrient limitation likely diminishes infectivity and disease dissemination, given that conidia serve as the primary inoculum. This reduction may also be linked to delayed melanization, as melanin positively influences conidiation in other pathogenic ascomycetes such as *Magnaporthe oryzae, Botrytis cinerea*, and *Aspergillus fumigatus* (20, 22, 23). Beyond sporulation, melanization contributes to host adhesion, appressorium formation, and overwintering survival, underscoring its multifaceted role in the disease cycle (23–25).

Carbon utilization assays revealed a pronounced preference for maltose, which supported melanization and sporulation at levels comparable to PDA. Given that PDA is rich in starch-derived carbohydrates, maltose likely reflects the carbohydrate environment encountered during host colonization. Maltose, a disaccharide generated during starch degradation, supported more robust development than glucose alone, suggesting that *N. dimidiatum* may be optimized for utilizing starch-derived sugars (26). Although sporulation was delayed relative to PDA, maltose was the only defined carbon source that supported both extensive melanization and appreciable conidiation. Similar maltose preferences have been documented in other plant-pathogenic fungi and ascomycetes, where uptake is mediated by dedicated transporters and intracellular hydrolysis via α-glucosidases (27). The ability of *N. dimidiatum* to grow on starch-containing media and secrete amylase further supports this hypothesis. Given that starch is present in pitaya tissues, these findings suggest that starch degradation and maltose assimilation may represent biologically relevant strategies during infection (28, 29).

In contrast, other sugars such as glucose and sucrose supported partial melanization but delayed or reduced sporulation, while polyols and lactose resulted in severely impaired growth. These differences likely reflect limitations in carbohydrate uptake or metabolic entry into glycolysis, as certain substrates do not readily feed into central carbon pathways (30, 31). Impaired glycolytic flux has been associated with reduced cyclic AMP-protein kinase A signaling, a pathway known to regulate melanin biosynthesis and fungal development (32), providing a plausible mechanistic explanation for the observed phenotypes.

Nitrogen utilization assays demonstrated a clear preference for complex, peptide-rich sources, with yeast extract supporting the most robust growth, melanization, and sporulation. Yeast extract contains peptides, amino acids, vitamins, and trace elements, which may collectively enhance fungal development compared to simpler nitrogen sources such as peptone or casein hydrolysate (9, 33). Although yeast extract does not fully mimic the nitrogen environment *in planta*, its efficient utilization suggests that readily assimilable nitrogen strongly influences fungal fitness. Melanization observed on ammonium-based sources indicates that *N. dimidiatum* can utilize inorganic nitrogen forms present in plant tissues; however, delayed sporulation under these conditions suggests that additional nutritional components absent from minimal media may be required for optimal development. The preference for peptide-rich sources aligns with the extracellular protease activity detected in this study, suggesting that host protein degradation contributes to nitrogen acquisition during infection (34).

Consistent with these metabolic preferences, we experimentally validated that *N. dimidiatum* secretes multiple extracellular enzymes, including cellulases, pectinases, amylases, and proteases. These findings corroborate genomic predictions of CAZymes and PCDWDE repertoires (9) and highlight their likely role in host colonization. Cellulose, starch, and pectin are major structural components of pitaya tissues, and their degradation facilitates nutrient acquisition and tissue colonization. Comparable enzymatic strategies have been described in other fungal plant pathogens, such as cutinases in *M. oryzae*, pectin lyase in *B. cinerea*, pectinases in *Sclerotinia sclerotiorum*, and endoglucanases in *Fusarium oxysporum* (35–38). Like *N. dimidiatum*, these pathogens exhibit necrotrophic stages during infection that necessitate breakdown of host macromolecules to sustain growth and disease progression. Expression of PCWDEs has also been linked to developmental processes such as conidiation, underscoring the interplay between enzymatic activity, pathogen fitness, and virulence (39). Extracellular protease may further support nitrogen acquisition and other infection-related processes, as reported for *A. nidulans* and *B. cinerea* (37–41). More broadly, the secretion of these enzymes is consistent with a virulence-associated strategy centered on host macromolecule degradation, suggesting that *N. dimidiatum* also secretes other proteinaceous factors, such as effectors or antimicrobial compounds, during infection (9, 42).

Taken together, this study establishes that *N. dimidiatum* exhibits distinct carbon and nitrogen utilization preferences and secretes a suite of extracellular enzymes capable of degrading plant-associated carbohydrates and proteins. Nutrient limitation significantly impairs melanization and sporulation, reinforcing the link between metabolic capacity and fungal fitness. While these findings provide a foundation for understanding the physiology and pathogenic potential of *N. dimidiatum*, future work should address carbon catabolite repression, host-derived nitrogen utilization, and the specific contributions of individual degradative enzymes during infection. Molecular and *in planta* studies will be essential for elucidating these mechanisms and developing effective strategies to mitigate dragon fruit canker.

## Author Contributions

J.F. conceived the study and designed the experiments. R.E.K. conducted the experiments. J.F., R.E.K., and R.G. wrote and revised the final manuscript.

## Acknowledgements

We thank colleagues whose work could not be cited due to space and scope limitations. We also thank the Fernandez laboratory for constructive feedback.

